# OmixLitMiner 2: Guided Literature Mining Tools for Automated Categorization of Marker Candidates in Omics Studies

**DOI:** 10.1101/2025.03.05.641607

**Authors:** Antonia Gocke, Bente Siebels, Jelena Navolić, Carla Reinbold, Julia E. Neumann, Stefan Kurtz, Hartmut Schlüter

## Abstract

Omics analyses are crucial for understanding molecular mechanisms in biological research. The vast quantity of detected biomolecules presents a significant challenge in identifying potential biomarkers. Traditional methods rely heavily on labor-intensive literature mining to extract meaningful insights from long lists of regulated candidates. To address this, we developed OmixLitMiner 2 to improve the efficiency of omics data interpretation, increase the speed for the validation of results and accelerate further evaluation based on the selection of marker candidates for subsequent experiments. The updated tool utilizes UniProt for synonym and protein name retrieval and employs the PubMed database as well as PubTator 3.0 for mining abstracts and full texts of available biomedical literature. It allows for advanced keyword-based searches and provides classification of proteins or genes with respect to their awareness level in relationship to scientific questions. OmixLitMiner 2 offers improved functionality over the previous version and comes with a user-friendly Google Colab interface. In comparison to the previous version OmixLitMiner 2 improves the retrieval and classification of relevant publications. The tool significantly reduces the time required for manual searches, as demonstrated in a case study involving proteomic data from spatially resolved mouse brain cortex layers.

**Statement of Significance:** We developed OmixLitMiner 2 to determine, for a given set of marker candidates, the extent to which they have been described in the literature. The tool is easy-to-use and can quickly generate a categorized list of references obtained by automated literature searches for protein and gene names and involving keywords related to a specific scientific question. The categorization provides a ranking of how well-studied the genes or proteins: Candidates of category 1 are well-known regarding the scientific question; Candidates of category 2 have been mentioned in a publication associated with the specific scientific question; Candidates of category 3 have never been described to be associated with the specific scientific question; Candidates of category 4 are not yet known proteins and are not associated with a gene name. This classification can aid in the decision, which protein or gene candidates to choose for follow-up experiments.

## Introduction

Proteomic, transcriptomic and genomic analyses have become an important part in the investigation of biological processes on the molecular level.^1^ They offer broad insights into the molecular mechanisms of the development of diseases. However, the vast number of molecules being quantified in these analyses, frequently exceeding several thousand biomolecules, present a significant challenge: identified proteins from this extensive dataset may already be well known, seldom mentioned or may have never been described with respect to the original scientific question^2^. Members of the latter two categories - not well known or unknown – could be potential new biomarkers or drug targets.

The identification of novel biomarkers or drug targets requires the discovery of candidates that exhibit a significant difference in their abundances between two compared phenotypes or a perturbed biological system and its control. Statistical tests usually identify hundreds of proteins or genes altered in their concentrations. In the past years, several pipelines for automated omics data analysis were developed, mainly focusing on creating protein-protein interaction networks and performing gene set enrichment analysis^3–5^. However, these analyses do not replace extensive literature mining for understanding the proteins’ specific function and association in biological processes in the investigated setting. Thus, the development of more literature mining tools could provide substantial benefits to the researchers.

The first crucial aspect in omics literature mining is the generation of synonym lists. Therefore, prior to undertaking literature searches in databases (e.g., PubMed^6^ or Google Scholar^7^), it is essential to compile a comprehensive list of all known gene synonyms. Furthermore, relevant and context-dependent keywords should be included to reduce the search space to the field of interest.

In response to these challenges, Steffen et. al. (2020) developed OmixLitMiner (OLM) as an automated tool for searching protein lists in PubMed.^8^ It employs a scoring method to classify candidates based on existing publications in conjunction with specific keywords into categories 1 (reviews available), category 2 (publication available) to category 3 (no publications). Because of modifications to the PubMed application programming interface (API), the initial version of the R-based OLM is no longer operational.

Recently NCBI released PubTator 3.0, a literature resource that employs artificial intelligence to search for genes and proteins in conjunction with diseases and other keywords within the PubMed literature database^9^. The tool provides more detailed results, as it integrates not only the keywords, but also new keywords with the same meaning as the original keywords, and facilitates the mining process for individual candidates. However, PubTator3.0 does neither enable simultaneous searches for multiple candidates nor allows for the classification of proteins based on their relevance and research status.

Thus, we present the new and improved tool OmixLitMiner2 (OLM2) to expedite the interpretation of omics data and thereby supporting the identification of potential biomarkers and drug target candidates. While OmixLitMiner retrieved a list of the gene synonyms, the OmixLitMiner 2 also retrieves and adds the protein names to the search queries to fully retrieve the available information of the investigated candidates enhancing the accuracy and comprehensiveness of the analysis. The literature mining tool still significantly reduces the time and effort typically required for manual searches of large number of marker candidates. This is especially aided by the integration of a new user-friendly Graphical User Interface within Google colab^10^, making the tool more accessible compared to the before available R package. Furthermore, the OmixLitMiner 2 also integrates the sophisticated literature mining capabilities of PubTator3.0.

## Material and Methods

### 1. Literature retrieval

#### a. OmixLitMiner 2

The OmixLitMiner 2 is composed of three major components: query building, query retrieval and categorization. It uses two major biomedical data collections: UniProt^12^ and the NCBI PubMed database^6^. The software is open source and well-structured as to simplify maintenance in case of updates such as new APIs. Unlike OmixLitMiner, OmixLitMiner 2 can be used via a Google colab notebook:

On the Google colab notebook one is guided by easy fillable forms to build the query. One needs to provide the protein list, specify whether the proteins are provided by their gene names or their accession numbers, give the taxonomy ID of the organism and the keywords, which represent the scientific question being investigated. A publication is selected only if it contains all the keywords.

To build the query, the Uniprot REST API is used to fetch the gene names, synonyms and now newly integrated the protein names for each protein. The user can choose whether the query should be answered by PubMed or PubTator3.

If the user chooses to use PubMed, the gene names, synonyms, and protein names are combined with the keywords provided to form the PubMed query. Additional features of the OmixLitMiner 2 include specifying if the tool should search for the keywords in only the title or in the title and the abstract, the maximum number of publications to be retrieved and lastly whether the publications should be ordered by publication date (from newest to oldest) or by their relevance (order used by the PubMed-API). The limit was set to 1,000, as the tool yields an Excel-file, which has limit of 1,048,576 rows and as to not restrict the number of proteins the tool can parse, as some combination can retrieve multiple thousands of publications, we restricted the number of publications being yielded from 10,000 to 1,000. This way multiple hundreds of proteins can be analyzed by the OmixLitMiner 2. The query is then supplied to the PubMed search API. The tool then parses the list of publications delivered and categorizes them according to the schema described below (Fig 4).

**Figure 1:**
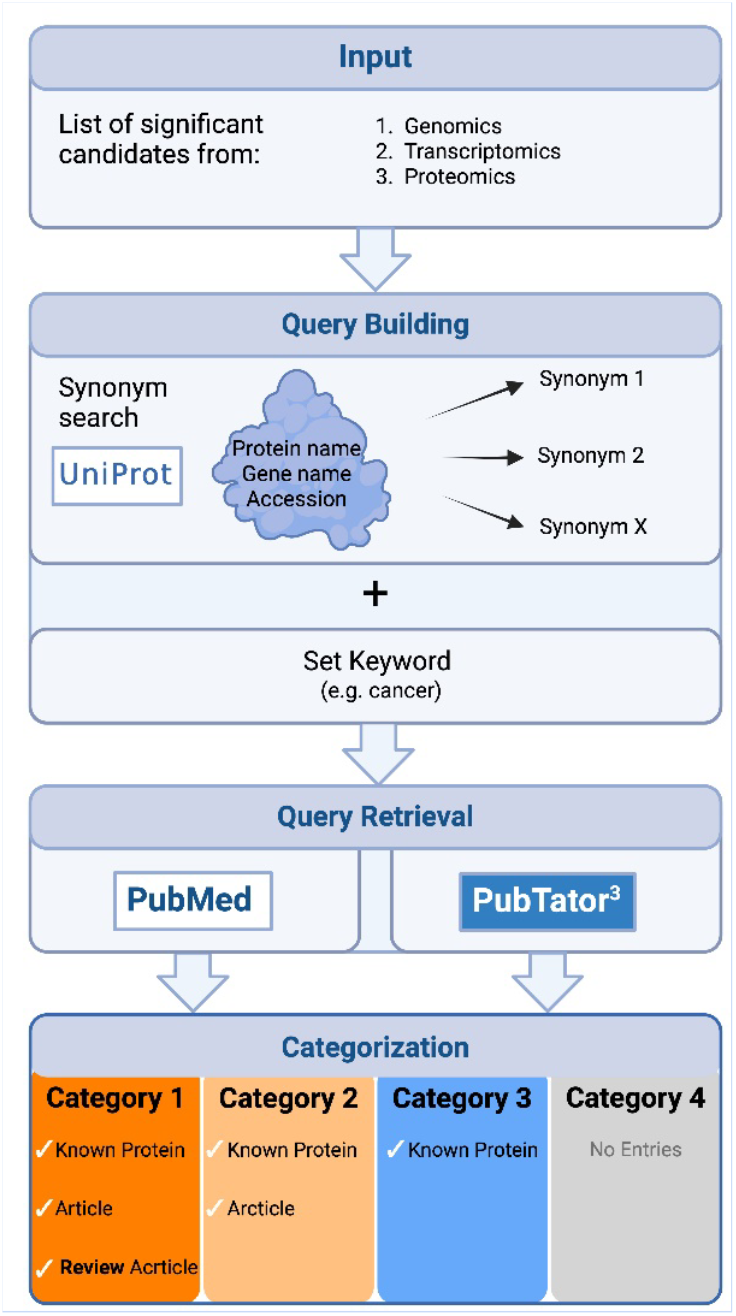
Overview of the literature mining workflow using OmixLitMiner 2.^11^.

**Figure 2:**
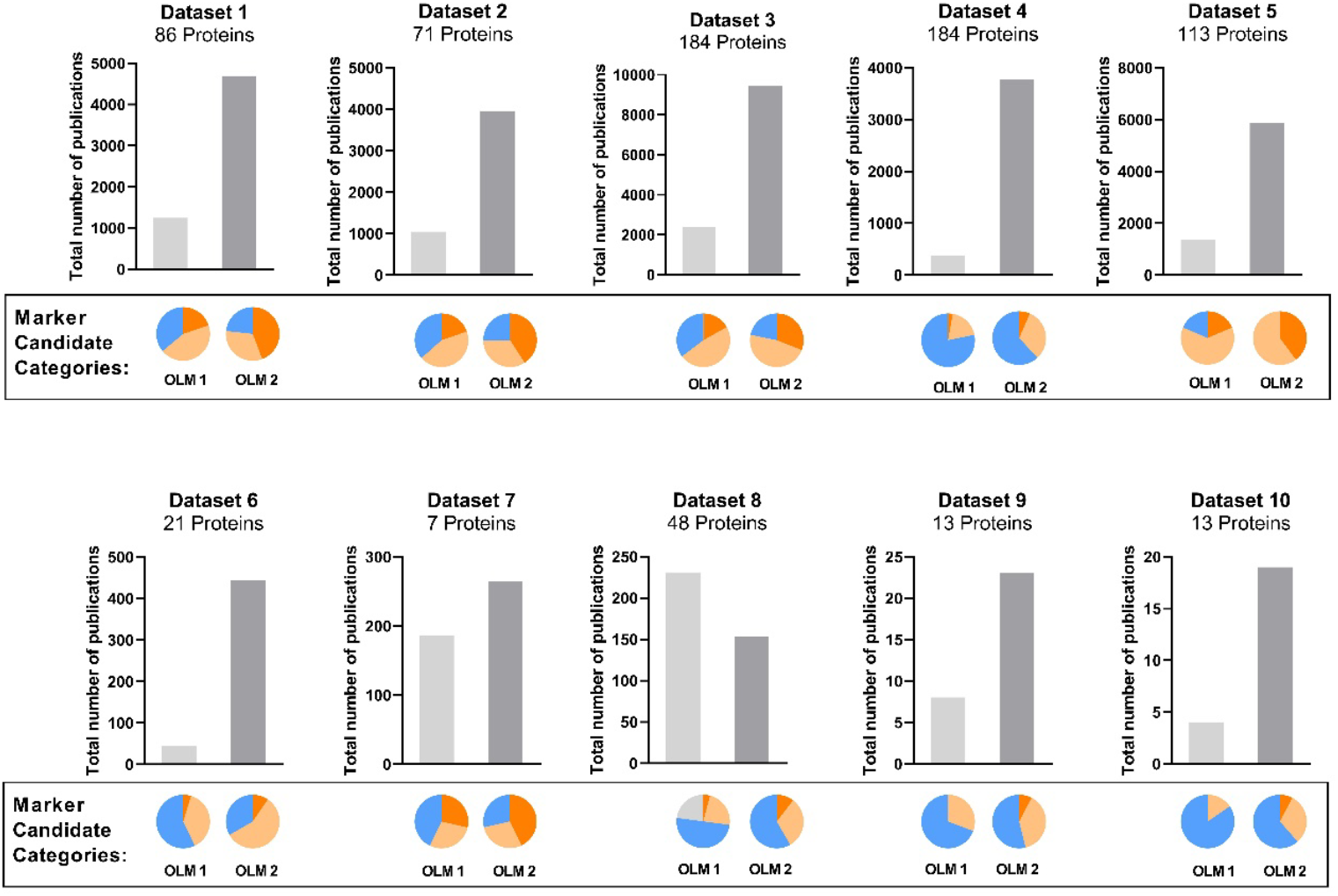
Comparison of OmixLitMiner 2 against the OmixLitMiner. Gene lists, key words and search results of the OLM were retrieved from the original publication by Steffen et al. Search results for OLM 2 were retrieved based on the same search setting including key words, genes limiting the search space to all papers published before or on 13.09.2019. For each gene a maximum of 1000 publications was set. In the plots, the sum of the number of publications for all gene names in the dataset is shown with the group distribution indicated in the pie chart above. Category 1: orange, 2: yellow, 3: blue, 4: grey. Dataset 1-5: Martinez-Aguilar et al.^19^, Dataset 6-7: Hänel et al. ^20^, Dataset 8: Mori et. al^21^, Dataset 9-10: Tian et al.^22^. Further dataset specifications can be found in supplementary table 1, detailed results can be found in Supplementary table 2-3).

**Figure 3:**
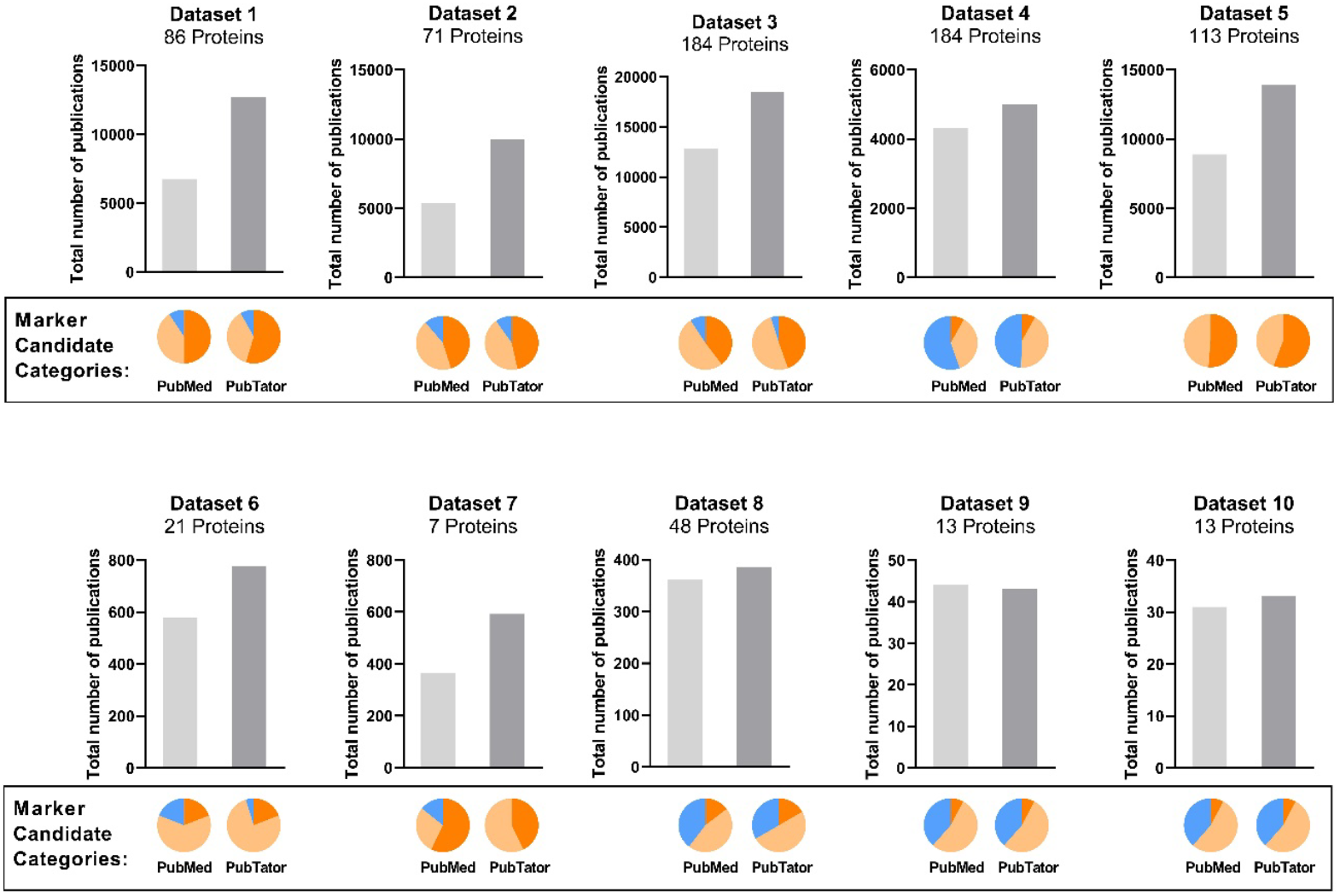
Comparing OmixLitMiner 2 performance using the PubMed and PubTator 3.0 API. For each gene a maximum of 1000 publications was set. In the plots, the sum of publications for all gene names in the dataset in shown with the group distribution indicated in the pie chart above. Category 1: orange, 2: yellow, 3: blue, 0: grey. Dataset 1-5: Martinez-Aguilar et al.^19^, Dataset 6-7: Hänel et al. ^20^, Dataset 8: Mori et. al^21^, Dataset 9-10: Tian et al.^22^. Further dataset specifications can be found in supplementary table 1, detailed results can be found in Supplementary table 4-5).

**Figure 4:**
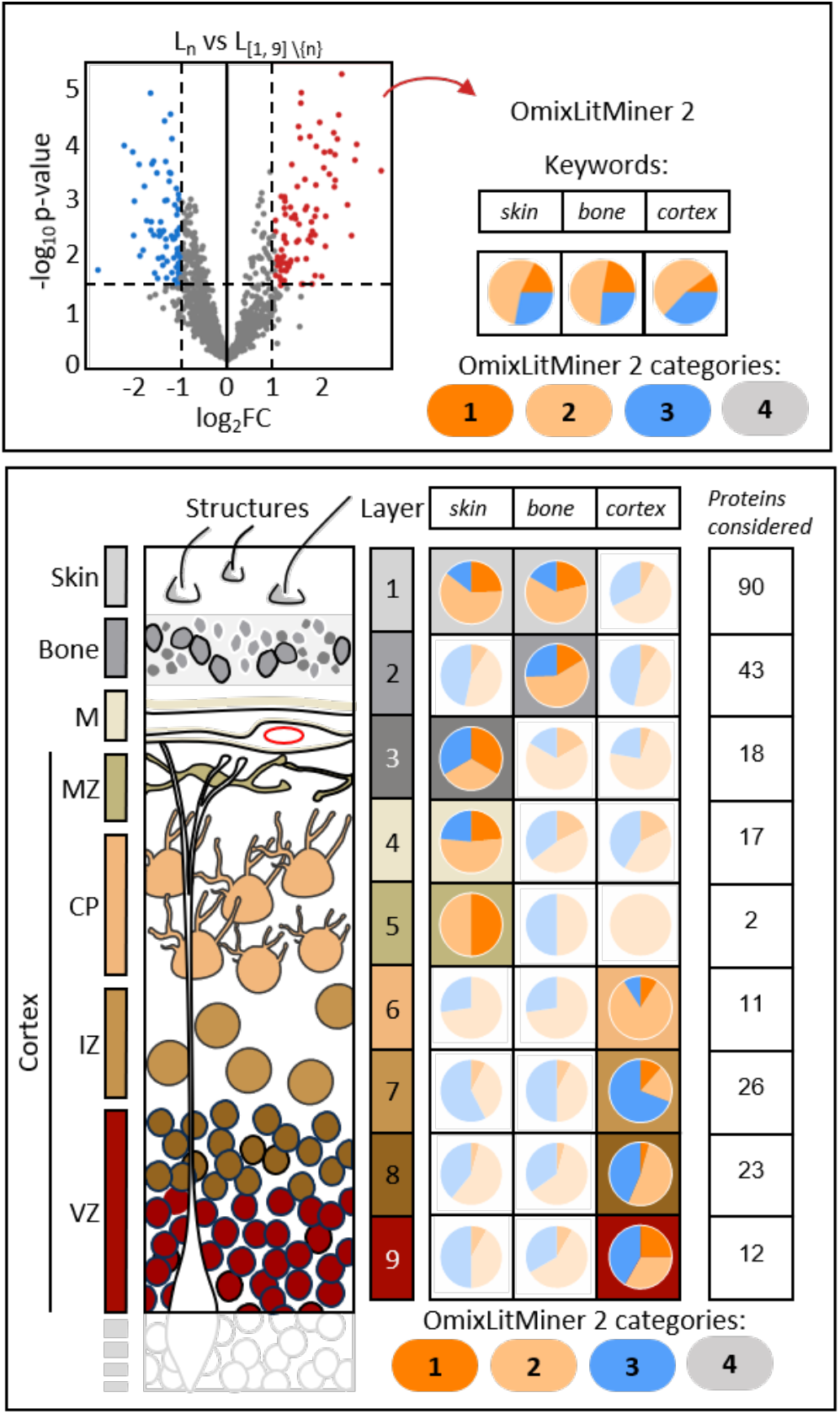
Literature Retrieval with OmixLitMiner 2to validate significantly higher abundant proteins for each layer. **Upper panel:** Significantly highly abundant proteins upon t-test (p < 0.05; log2FC > 1) comparing a single layer against the rest of the layers (Ln vs L[1, 9] \{n}) were selected. These proteins were assigned according to the keyword: skin, bone or cortex. The output is displayed as a pie chart labeled with the number of proteins used in the search. Lower panel left: Shows the anatomical structures of the ablated areas from the skin surface to the bone, meninges into the cortex. M = meninges, MZ = marginal zone, CP = cortical plate, IZ = intermediate zone, VZ = ventricular zone. Lower panel right: Shows for each layer 1 to 9 the respective pie plot of the OLM2 result with the highest category 1 proportion.

If the user chooses to utilize the PubTator3 API, the marker candidates are pre-processed analogous to the PubMed, except for the interaction with the different API. Additionally, no limit is set for the number of publications by the authors, as the PubTator search API only retrieves the first 10 publications, but links to all search results are provided in the output file.

### 2. Categorizing

The OmixLitMiner 2 categorizes the genes/proteins based on the literature retrieval described in the previous section. If, for a given protein P, the literature retrieval with the query build for P delivers at least one article/publication, P is assigned to either category 1 or 2. P is assigned to category 1 if one of these publications is a review article. If no publication was delivered by the literature retrieval with the query build for P, but P is a valid identifier for a protein in UniProt, P is assigned to category 3. In all other cases, P is assigned to category 4.

### 3. Validation

While there are literature mining tools, which help to retrieve information about specific genes, not many literature mining tools automate the retrieval of publications for multiple genes simultaneously. Additionally, at the time of writing, the authors are not aware of any literature mining tool that integrates large lists of possible markers and keywords for marker classification. While GenCLiP 3^13^ also uses large gene lists and certain keywords as an input, however instead of classifying the genes, it generates co-occurrence networks for these genes, indicating possible interactions. Similarly, GIREM^14^ also focuses more on reconstructing co-occurrence networks and integrates the semantic relationship between the co-occurring genes to generate more accurate networks.

Thus, other than the OmixLitMiner, there was no other suitable tool against which we could validate our results.

#### 3.1. Comparison of OmixLitMiner and OmixLitMiner 2

As the API of PubMed has been updated, we cannot compare the OmixLitMiner 2 directly against the OmixLitMiner R package. However the authors of the OmixLitMiner published several test data sets, which could be reprocessed with the OmxLitMiner 2 by setting a maximum date to the date of retrieval of the original OLM publication (2019-09-13)^8^. Since the publication of the OmixLitMiner multiple Accession numbers have been deprecated, thus we mapped the accession number to gene names and performed the search with these. Further dataset specifications can be found in supplementary table 1.

#### 3.2. Functional Validation and Application Example

The data we used for a functional validation and an application example was published by Navolić et al., 2023^15^. The harmonized data (Table S2) includes spatially resolved proteome data of brains from wild type mice at embryonal day 14 and processed as described by the authors.

Proteins with 70 % valid values were used for t-test comparison (limma version 3.54.2) for each layer against the other layers (L_n_ vs L_[1, 9] \{n}_). The significantly high abundant proteins (p < 0.05; log_2_FC > 1) were selected (SI Appendix Table 1) and categorized using the OmixLitMiner 2 combined with single keywords covering the main anatomical structures: skin, bone, and cortex. The PubMed API was used and the search was restricted to the title and abstract. The proportion of proteins for each category was displayed as a pie plot for the respective layer. The pie plots with the highest number of Category 1 proteins were highlighted by a different colored background (Figure 5).

**Fig. 5:**
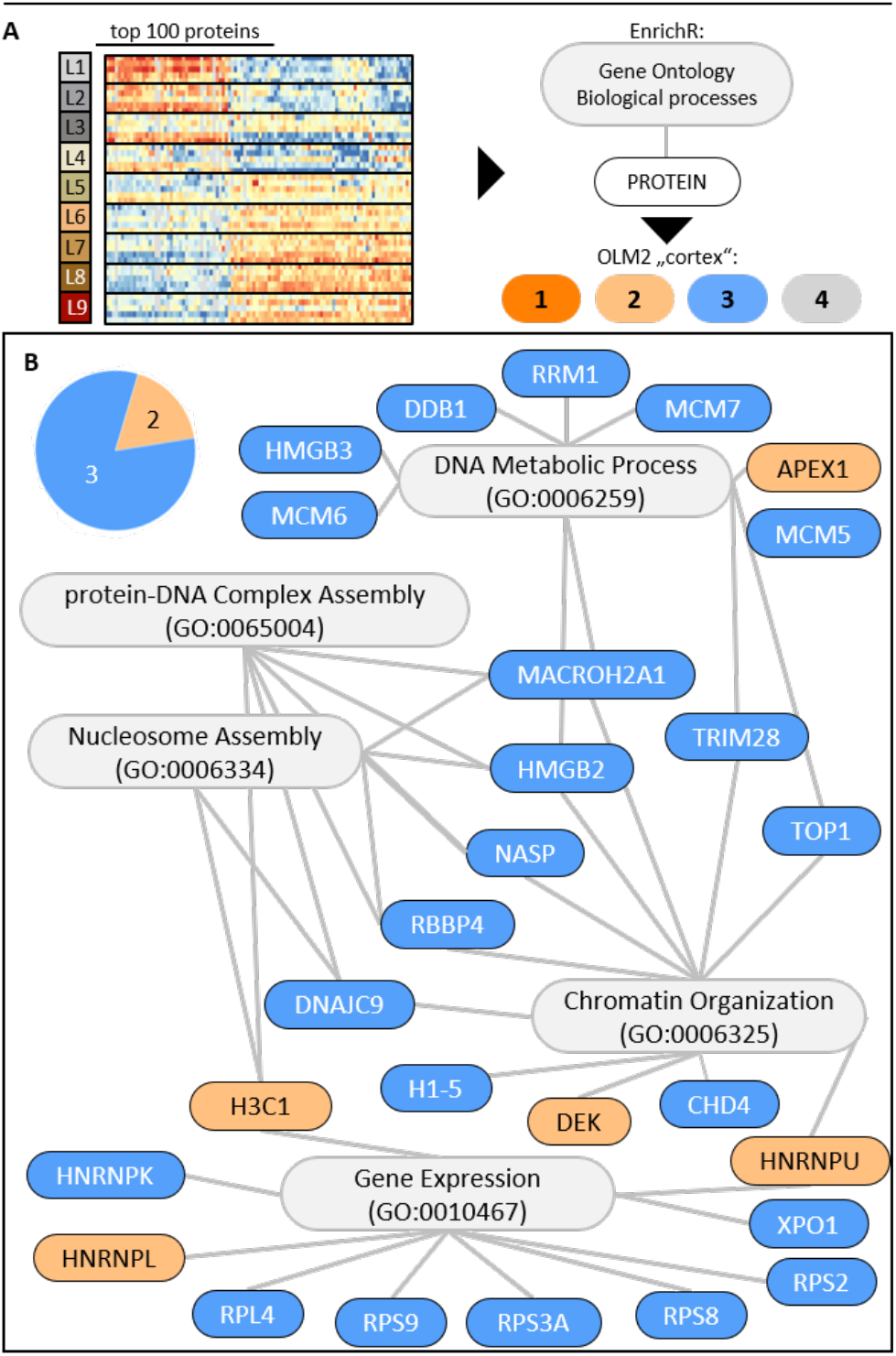
Categorization of gene ontology associated proteins. Upper panel: Top 100 significant proteins based on ANOVA across layer 1 to 9^15^. These proteins were analyzed using gene ontology for biological processes (GO:BP) in Enrichr^16–18^. Lower panel: Top 5 GO:BP-terms and their related proteins are displayed. Each protein is colored according the category assigned by OLM 2 when searching with the keyword *cortex*.

The top 100 proteins based on ANOVA between all layers were obtained from Navolić et al., 2023^15^ (Supplementary table 6). These proteins were used to determine their function involved in the gene ontology (GO) for biological processes (GO:BP) with Enrichr ^16,17,18^. The proteins associated with the top 5 GO:BP-terms were further categorized using OmixLitMiner 2 with keyword = cortex, the PubTator API and KeywordInTitleOnly = TRUE. The terms and related proteins are plotted as a bipartite graph (igraph version 2.0.3) and the proteins are colored according to the category they are assigned to.

## Results and Discussion

### OmixLitMiner and OmixLitMiner2: Comparison of new functions

The new OmixLitMiner 2 offers multiple benefits and improvements compared to the OmixLitMiner. **Table 1** lists the main features of the both software tools tools.

**Table 1:** Main features of OmixLitMiner and OmixLitMiner2.

|  | OmixLitMiner | OmixLitMiner 2 |
| --- | --- | --- |
| **Implementation language** | R | Python |
| **Database** | PubMed | PubMed<br>PubTator 3.0 |
| **Searches** | Title + Abstract | Title + Abstract |
| **Keyword Combination** | AND | PubMed: AND<br>PubTator: AND and with similar words* |
| **Input** | Accession number or gene name | Accession number, gene name or protein name |
| **Maximum number of papers** | 1000 | PubMed: 1000<br>PubTator 3.0: No limit (via link, only 10 retrieved for output table) |
| **Accessibility** | R package (deprecated) | Google Colab + GitHub |
| **Categories** | 0, 1, 2, 3 | 1, 2, 3, 4 |

\*Through large language model of PubTator 3.0

### Comparison of search results from OmixLitMiner and OmixLitMiner 2

To compare the performance of OmixLitMiner 2, we chose the same 10 datasets used in the original publication comprising proteomics data^19–21^ and genomics data^22^. The datasets are comprised of different previously published data of different biological areas of interest. As a result, for each data set different keywords were used (e.g., *Cancer, Metastasis, Thyroid, Migration*).

The result of the search shows that OmixLitMiner 2 retrieved significantly higher numbers of publications for 9 out of 10 datasets (**Figure 2**). Moreover, the proportions of protein-to-category assignments changed. For dataset 1 containing 86 candidates, the number of proteins assigned to category 1 increased from 17 to 38. Overall higher numbers of proteins in category 1 could be found for all datasets when using OmixLitMiner 2. For all datasets, except dataset 8, OmixLitMiner 2 assigned fewer proteins to category 3. The exception for dataset 8 can be explained by the fact that with OmixLitMiner 2, the number of proteins of category 4 (former category 0) candidates was reduced from 11 to 0. Moreover, the high number of publications found by OLM is caused by an erroneous output when proteins were assigned to category 0 but still 11 publications related to the keyword “metastasis” were mapped. With OLM 2, for all candidates, successful synonym search on UniProt was now conducted, as no category 0 appears and no wrong papers were retrieved.

Especially for the datasets derived from genomics data (9 and 10), the number of publications and the retrieval of category 2 and 1 candidates in combination with the corresponding keyword increased significantly. This is due to the inclusion of protein names into the literature mining.

### Comparison of PubMed and PubTator 3.0-based searches

With the PubTator 3.0 AI-based literature mining the sensitivity for the keywords is increased by incorporating similar and related words. For all datasets, except Dataset 9, PubTator 3.0-based search retrieved more publications than PubMed-based search. Also, the number of category 1 proteins (review article found) increased significantly, highlighting the benefit of Pubtator 3 based keyword search. For Dataset 9, 1 more publication was found for the candidate MEN1, also known as Menin, when using the PubMed-based search. However, this publication focusses on MEN-1, multiple endocrine neoplasia 1 (MEN-1) syndrome, and not on our candidate. Even though the request of the PubMed API did not contain the name MEN-1, the publication was retrieved. In contrast, PubTator 3.0 did not retrieve the wrong publication, highlighting the benefits of using the large language model-based search.

### Application examples of OmixLitMiner 2 in research

As an application example we use OmixLitMiner 2 to validate morphologic layers of an embryonic mouse head after proteomics analysis based on data published by Navolić et al.^15^. As depicted in **Figure 4**, 9 layers were ablated and analyzed via quantitative differential LC-MS/MS-based proteomics comparing each of the layers against all 8 other layers. Several significantly higher and lower abundant proteins between the different layers of the cerebral cortex were identified, as visualized in the volcano plot. We used OmixLitMiner 2 in a search previously described marker proteins to retrieve literature allowing to validate the different layers.

In particular, the significant proteins between the different layers were categorized based on their association with the specific keywords *skin, bone*, and *cortex*.

These categorizations are visualized in the pie charts, where each section represents the proportion of proteins assigned to categories 1, 2, or 3.

For Layer 1, significantly differential proteins were predominantly assigned to category 1 and 2 when searching with the keyword *skin*, indicating marker proteins for the skin and thus, validating these layers as layers including the skin. For layer 2-4, review articles (Category 1) were found for all the three keywords *skin, bone* and *cortex*, while more review articles were found for *skin* and *bone* indicating the layers belong to the skin, bone and meninges of the brain. However, the highest number of category 1 papers were identified in the skin and bone searches. For layer 5, only 2 marker proteins were available, thus limiting the search space. For layer 6-9 mostly review articles were found for the keyword *cortex* validating the layer assignment to the cortex of the brain.

This analysis revealed the nuanced distribution and existing literature support for proteins across different tissue layers, demonstrating the efficacy of OmixLitMiner 2 in facilitating targeted literature mining and functional categorization of proteomic data.

While the validation process is used to assign the potential markers to already reviewed literature, a second use case of the OmixLitMiner 2 considers proteins assigned to category 3 as potentially new marker proteins.

In our second use case, gene set enrichment analysis using Enrichr based on “Gene Ontology: Biological Process”, was performed on the top 100 ANOVA-significant proteins across all 9 layers of the dataset from Navolić et al.^15^ The proteins were assigned to different gene sets, namely DNA Metabolic Process, Chromatin Organization, Gene Expression Nucleosome Assembly and protein-DNA Complex Assembly (**Figure 5)**. With OLM 2 it is now possible to add a second layer of classification to the proteins. When using OLM 2 with the keyword *cortex*, proteins assigned to category 2 or 3 are not as well known in the literature. If they belong to a particular biological process of interest, they may be interesting candidates for further analysis. The integration of such additional information from enrichment analyses has the potential to reduce the number of targets that need to be investigated.

## Conclusion

Here we present an improved literature mining tool, OmixLitMiner 2, which aims to aid researchers in sorting and classifying a large number of genes or proteins from omics analyses regarding specific scientific questions. We benchmarked the tools against the previous version, presenting an overall improvement in the number of retrieved publications and in the sensitivity of the categorization. Finally, we present two approaches for utilizing the OmixLitMiner 2 results. The first approach employs OmixLitMiner 2 for the validation of results obtained by other methods, while the second approach shows how researchers can further reduce the number of potential proteins to investigate by combining the results of the OLM 2 with Gene Ontology information with the results of the OmixLitMiner 2. Both approaches aim to help researchers in streamlining their work and research process.

## Data availability

The OmixLitMiner 2 can be accessed and used via: https://colab.research.google.com/drive/17FJ4kBt1iSBwqk5vTuIWvDqcB3UVCrCC?usp=sh aring

## Funding Information

### Author Contributions

Conceptualization, H.S., B.S.;

software OLM2, A.G., C.R.;

software testing, A.G., B.S.;

validation, A.G., J.N.;

benchmarking, A.G.;

case study, J.N.;

data curation, A.G., B.S.;

writing—original draft preparation, A.G., B.S., J.N.;

writing—review and editing, A.G., B.S., J.N., J.E.N, S.K., H.S.;

visualization, J.N.; B.S.; A.G.,

supervision, H.S., S.K., J.E.N.;

project administration, B.S., H.S.;

funding acquisition, H.S.

